# On the Importance of Polarizability in Simulations of 2D IR Spectra of the Potassium Ion Channel KcsA

**DOI:** 10.64898/2026.11.20.768138

**Authors:** Rasanjali L. Ranawaka, Alexei A. Kananenka

## Abstract

The bacterial KcsA channel is a K^+^ channel which is functionally homologous to mammalian ion channels. It assembles as a cone-shaped tetramer that is embedded in the membrane. Its narrowest conserved region, known as the selectivity filter (SF) is comprised of five distinct ion-binding sites (S0–S4) that coordinate the selective permeation of K^+^ ions. Recent two-dimensional infrared (2D IR) spectroscopy experiments, together with molecular dynamics and line shape simulations on the isotope-labeled SF residues V76 and G77, suggest that in the closed conductive state either S1/S3 or S2/S4 are occupied by ions with water molecules in between, consistent with the soft knock-on mechanism. 2D IR spectroscopy probes amide I vibrations that are highly sensitive to local environmental fluctuations thus providing a powerful approach for distinguishing ion permeation mechanisms. However, current 2D IR spectral calculations are largely restricted to non-polarizable force fields, which may not fully capture the realistic channel behavior. Here, we report 2D IR simulations based on all-atom molecular dynamics trajectories of the KcsA channel embedded in a lipid membrane performed using nonpolarizable CHARMM36m, polarizable CHARMM-Drude, and CHARMM force field with electronic continuum correction. The spectra were calculated for V76 and G77 isotope-labeled SF and for soft knock-on configurations. Our results show that simulations with the polarizable CHARMM-Drude force field better reproduce experimental 2D IR spectra underscoring the importance of including the polarization effects in simulations of ion channels.

## 1 Introduction

Potassium (K^+^) ion channels are transmembrane proteins found in neurons, muscle, and cardiac cells [1–3]. They play a crucial role in cell biology, primarily in maintaining the resting potential of the membrane, shaping action potentials, and regulating cell excitability. Malfunctioning K^+^ ion channels lead to disorders such as potassium channelopathies [4]. Therefore it is important to understand the structure and function of an ion channel at an atomistic level for effective treatment [5, 6].

The KcsA channel, derived from the bacterium *Streptomyces lividans*, is one of the most widely studied K^+^ channel due to its high structural and functional similarity to mammalian potassium channels [7–9]. KcsA is a cone-shaped tetramer with a 3 Å wide narrow region [10] called the selectivity filter (SF) which is composed of the signature pore-lining residues: T75-V76-G77-Y78-G79 (TVGYG) [11, 12]. The backbone carbonyls of these residues, along with the side-chain hydroxyl group of T75 residue, form five ion-binding sites, S0-S4, from the extracellular to intracellular side [7, 13–16], as shown in Figure 1A. The SF sequence endows the channel with rapid ion conduction because of its high selectivity for K^+^ over smaller ions such as Na^+^ [13, 17–20].

**Figure 1:**
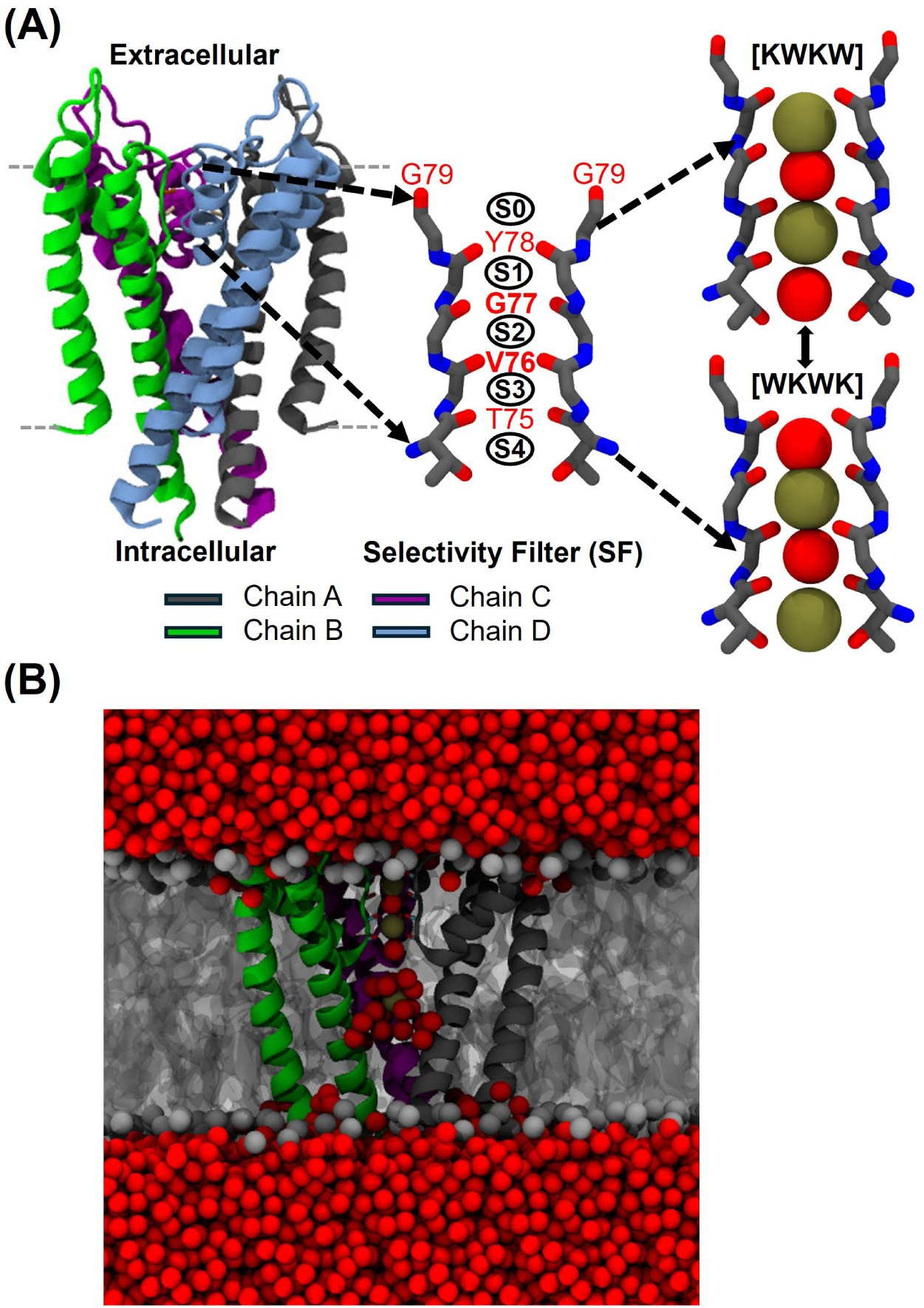
(A) The KcsA potassium channel (left), the selectivity filter (center), and the two soft-knock configurations used in this work (right). Selectivity filter residues V76 and G77 for which 2D IR spectra are calculated are highlighted in bold. (B) The simulation system showing the KcsA channel embedded in a DPPC lipid membrane with explicit water and ions. The K^+^ ions and water molecules are shown in olive green, and red, respectively.

The X-ray crystallographic study of the KcsA channel by Zhou *et al.* [21] indicated that the SF holds an average of two K^+^ ions, alternating between S1 and S3 or S2 and S4 sites, with water molecules intervening to buffer ion-ion electrostatic repulsion [21, 22]. Further the alternating ion/water SF sequence was confirmed by electrophysiological single-channel recordings and, more recently, by two-dimensional infrared (2D IR) spectroscopy [22–24]. The latter experiments were interpreted in terms of a mixture of two configurations which we denote as [KWKW] and [WKWK], where K and W represent the K^+^ ion and water molecule, respectively, occupying sites S1-S4 as shown in Figure 1A [22–24]. The SF fluctuate between these two configurations until a third K^+^ ion approaches the intracellular entrance of the SF and shifts the single file of ions and water up causing an S1 ion to leave the SF.

Alternatively, X-ray crystallography and nuclear magnetic resonance (NMR) experiments suggest that two ions occupying adjacent binding sites drive the ion transport [25–29]. This mechanism has come to be known as direct knock-on (or hard knock) [26, 30]. Studies reveal that S2 and S3 sites are frequently occupied by K^+^ ions simultaneously in the hard-knock mechanism suggesting a [WKK0] might be a prevailing configuration (“0” represents an empty binding site) [25–27, 31, 32]. Despite a wealth of experimental and simulation data [14, 16, 21, 22,24,25,33–37] the precise ion transport mechanism and its dependence on the conditions such as transmembrane potential and ion concentration is still not fully known [28].

2D IR spectroscopy is an ultrafast technique that probes molecular vibrations [33, 38–40]. The most extensively studied vibration in proteins is the amide I mode, which arises primarily from carbonyl stretching vibrations of the protein backbone [41, 42]. The amide I vibrational frequency is a sensitive probe of the local electrostatic environment which was utilized to study hydrogen bonding, solvent exposure, and ion binding [41, 43–45]. Site-specificity of 2D IR spectroscopy can be achieved using residue-specific isotope labeling [41, 43–48]. Labeling naturally more abundant ^12^C^16^O to ^13^C^18^O redshifts the vibrational frequency by 66 cm*^−^*^1^ [49, 50] allowing to routinely resolve the signal from the labeled site from the rest of the amide I modes of the protein. This approach was leveraged in recent 2D IR experiments on the KcsA channel [24, 33, 35].

Often 2D IR spectra are interpreted with the help of molecular dynamics (MD) simulations and line shape modeling [24, 33–35, 51]. Ryan *et al.* [24], following earlier simulations by Strong *et al.* [34], in two independent experiments measured 2D IR spectra of the KcsA channel with isotopically labeled carbonyl groups of V76 and G77 residues. Simulated spectra clearly showed the prevalence of the soft-knock ion configurations at the equilibrium conditions of the experiment [24]. In simulations, local electric fields due to solvent, ions, lipids, and protein side-chain atoms, are related to amide I vibrational frequencies using the so-called spectroscopic maps [51, 52] allowing to map spectral features to the underlying structures.

2D IR simulations are based on atomistic configurations generated in MD simulations. In Refs. [23, 24, 34, 35] 2D IR spectra were calculated from MD trajectories generated using GRO-MOS96 53a6 force field [53–55]. 2D IR simulations carried out under applied transmembrane potentials by Kopec *et al.* [25] employed amber99sb force field [25,56] and supported hard-knock configurations.

GROMOS and amber, together with another popular force field CHARMM [57] belong to a family of nonpolarizable force fields. A known limitation of non-polarizable force fields is their inability to explicitly model induced polarization. Consequently, they are unable to accurately describe local microscopic interactions and thermodynamic properties of ions; frequently overestimating electrostatic repulsion and free-energy barriers within low-dielectric, confined environments such as SF [58].

Studies demonstrate that explicit electronic polarization is important for accurately describing the stability and dynamics of buried ion pairs and the hydration of protein cavities [59, 60]. The two most widely used polarizable force fields are CHARMM-Drude [61] and AMOEBA [62], in which the polarizability is computed through Drude oscillator [63–65] and induced dipole models [66, 67], respectively. CHARMM-Drude simulations of ion channels have a long history [61, 68–71]. It has been shown that CHARMM-Drude model improves the prediction of ion-pair stability, captures greater structural fluctuations, and better reproduces water positions and ion occupancy [59, 60]. In a study of hydrophobic gating in Big K^+^ channels with the CHARMM-Drude force field an enhanced gating efficiency has been demonstrated [61]. Simulations based on AMOEBA force field elucidated a critical role of polarization in K^+^ ion channels and ion binding thermodynamics [62]. Comparative simulations using AMOEBA and CHARMM-Drude polarizable force fields, alongside nonpolarizable CHARMM models, have shown that, although hydrophobic gating remains qualitatively consistent across models, the treatment of polarization can influence pore hydration, ion–protein interactions, and ion-permeation energetics in ion channels [72, 73].

The electronic continuum correction (ECC) is an alternative approach that introduces polarizability implicitly by scaling the atomic charges [74–78]. CHARMM force field with ECC scaled charges has been recently used to calculate various properties of NaK2K, TRAAK, and MthK channels [79]. The calculations with ion charge scaled by 0.78 reproduced experimental current-voltage (*I*–*V*) curves. Equilibrium molecular dynamics simulations at zero membrane potential show MthK channel conductances of only 5–9 pS at 400 mM KCl across AMBER and Drude force fields, representing a 30–60-fold underestimation relative to experiment that underscores fundamental inaccuracies in current force field representations, their underlying hard-knock mechanisms, and the premise that scaling ionic charges *via* the ECC offers a viable solution [37]. Evaluating 29 force fields on the MthK potassium channel, Uriarte *et al.* [69] further showed that SF K^+^ and water occupancy is highly sensitive to delicate balances in ion–water, ion–backbone, and water–backbone coupling, with three force fields supporting the soft knock-on and the four force fields supporting hard-knock mechanisms, while 22 yielded intermediate occupancy states.

In this Article we critically assess the importance of including polarization in modeling 2D IR spectra of SF. We consider a three-tier approach which includes a non-polarizable force field CHARMM36m, an ECC modified CHARMM force field which includes polarization in a mean-field way, and CHARMM-Drude force field which treats polarization explicitly. 2D IR spectra of isotope-labeled V76 and G77 residues were calculated based on MD trajectories of [KWKW] and [WKWK] configurations generated using the three different types of force fields. The accuracy of simulated 2D IR spectra was assessed by comparison with the experimental 2D IR spectra of Ryan *et al.* [24].

## 2 Materials and Methods

### 2.1 Molecular Dynamics Simulations

Two systems corresponding to the soft knock-on (Figure1A) configurations were built using the wild-type KcsA channel (PDB ID 1K4C [7], residues 22–124) using CHARMM-GUI platform [80]. The GalaxyFill algorithm in CHARMM-GUI [81] was employed to model the incomplete side chains of channel residues, which were protonated at pH 7.5 with neutral GLU, the protonation is important for KcsA stability [82]. The N- and C-terminus were modeled using the CHARMM NTER (–NH_3_^+^) and CTER (–COO ^−^) patches, respectively, to represent their physiologically charged states of the free terminal residues at neutral pH. The crystallographic water molecules and K^+^ ions were preserved, except for those within the SF, where some ions were replaced with water molecules, using Visual Molecular Dynamics (VMD), [83, 84] as required to generate the desired ion configurations. The resulting structures were embedded in Dipalmitoylphosphatidylcholine (DPPC) bilayers and solvated with TIP3P [85] water containing 500 mM KCl, mimicking experimental conditions with high local ion concentrations. The final system, comprising 73,146 atoms in a box with the dimensions 93 *×* 93 *×* 92 Å^3^, was subjected to sequential energy minimization and equilibration using the CHARMM36m [57] non-polarizable force field, following the standard CHARMM-GUI protocol described below. The equilibrated structures were then used to construct CHARMM-Drude force field systems with the Drude-Prepper [86] input generator module in CHARMM-GUI, solvated in an SWM4-NDP [87] water box (95 *×* 95 *×* 87 Å^3^). The final system contained 116,002 atoms and Drude particles.

#### 2.1.1 Simulations with CHARMM36m force field

The energy minimization with the CHARMM36m force field was performed for 3,000 steps using the NAMD 2.14 software [88]. Scaled positional restraints were applied to the protein backbone (10.0 kcal/mol/Å^2^) and side chain heavy atoms (5 kcal/mol/Å^2^), as well as to the DPPC lipid head groups (5 kcal/mol/Å^2^) and dihedrals (500 kcal/mol/Å^2^). The system was then thermalized to 298.15 K in the NVT ensemble for 125 ps with a 1 fs timestep. Constraints were gradually relaxed over another 125 ps before switching to NPT dynamics for an additional 125 ps (1 fs timestep). During this stage, the cavity ion was restrained with a 5 kcal/mol/Å^2^ force constant, which was maintained throughout the simulation. Subsequently, relaxation dynamics were performed for 1.5 ns in the NPT ensemble using a 2 fs timestep, with the SF–C*α* restraints reduced to 2 kcal/mol/Å^2^. During NPT equilibration, the ions in the SF were softly restrained with a 1 kcal/mol/Å^2^ force constant. The equilibrated structures served as the initial conformations for the Drude simulations. Production simulations were then carried out for 4.5 ns using flat-bottom half-harmonic potential restraints on the K^+^ ions (similar to the approach used in Ref. [89]) within the SF, together with the SF–C*α* and cavity ion restraints, as part of the equilibration process to prevent significant ion shifts. During the production simulation, the trajectory frames were saved every 20 fs for 2D IR spectral simulations. In what follows we will refer to CHARMM36m as CHARMM for brevity.

#### 2.1.2 Simulations with CHARMM-Drude force field

The Drude-particle-based system was energy-minimized for 200 steps, followed by NPT equilibration for 100 ps with a 0.5 fs timestep. Production simulations were performed in NVT ensemble for 2 ps with a 1 fs timestep. During the production simulation, the trajectory frames were saved every 20 fs. Throughout the production phase of the Drude simulations, positional restraints (5 kcal/mol/Å^2^) were applied to all protein C*α* atoms except those in the SF (residues 74–79) to avoid significant deviations from the starting configuration [23]. Each of the 100 frames generated as described above was used as starting structures for 20 ps trajectory for 2D IR calculations. During the 20 ps simulations only the C*α* atoms of the SF residues were constrained with the same force constant [23]. Flat-bottom half-harmonic potentials were also applied to the ions in the channel, consistent with earlier stages, to preserve ion configurations.

#### 2.1.3 Simulations with CHARMM-ECC force field

The same initial configurations and protocol used for the CHARMM simulations were used to simulate a system in which the ion charges were scaled according to the ECC approach and following Hui *et al.* [79]. In their study of potassium ion channels, Hui *et al.* demonstrated that scaling the ion charges (K^+^/Cl^−^) in the system by a factor of 0.78, together with the radius-corrected Lennard-Jones parameters, effectively accounts for electronic polarization, and allowed them to reproduce experimental I-V curves and ion selectivity [79]. To maintain the system’s overall charge neutrality, the charges of all side chains atoms of ASP, HSD, PRO, GLU, and ARG residues were also scaled as well as the charges of NTER and CTER patches. Radius-corrected Lennard-Jones parameters were taken from Ref. [79]. The equilibration and production simulations were carried out exactly as the CHARMM simulations described above.

### 2.2 2D IR spectral calculation

2D IR spectra were calculated from CHARMM, CHARMM-ECC, and CHARMM-Drude trajectories generated as described above. It should be noted that, because only the ion and protein C*_α_* positions were restrained, the water molecule occupying the extracellular S1 site was found to exit the SF and then re-enter, causing the S1 site occupied only intermittently. To ensure the structural consistency in the SF ion/water occupancy across CHARMM, CHARMM-ECC, and CHARMM-Drude spectral calculations, equilibrated CHARMM and CHARMM-ECC structures were selected based on their similarity to the equilibrated CHARMM-Drude ion/water configurations. The RMSD values were calculated by aligning the carbonyl atoms of the SF residues (residues 75-79) to the equilibrated CHARMM-Drude reference structures. The resulting structures, exhibiting RMSD values between 0.21 and 0.88 Å relative to CHARMM-Drude were used for continuous CHARMM and CHARMM-ECC spectral calculations. This approach isolates force field effects from structural variations, as only the former are the focus of this study. All 2D IR calculations used 100 20 ps trajectories with the system configurations saved every 20 fs. 2D IR spectra were simulated for V76 and G77 ^13^C^18^O isotope labeled residues following experimental work of Ryan *et al.* [24, 35].

2D IR spectra were calculated using a hybrid quantum-classical approach [51, 90, 91]. The vibrational frequency of the *i*th isotope labeled amide I group was determined as the sum of electrostatic and nearest-neighbor frequency shifts as well as the frequency shift due to isotope labeling

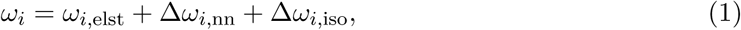

where the first term, the electrostatic frequency map, describes the amide I vibrational frequency (in cm*^−^*^1^) as a function of the local electric field and was taken from Ref. [52]

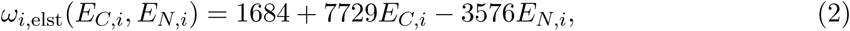

where *E_C,i_* and *E_N,i_* are the electric fields (in atomic units) calculated at the positions of the *i*th amide C and N atoms and projected onto the amide C=O bond, respectively. Following the original work of Wang *et al.* [52] the electric fields were calculated from all protein atoms that are more than three covalent bonds away from the amide I group, and all other atoms including water, ions, and lipids. Only atoms with within a cutoff of 20 Å from the C or N atom were included in the calculation. In the CHARMM calculations the original (CHARMM36m) atomic point charges were used. In CHARMM-ECC calculations the scaled atomic charges described in Sec. 2.1.3 were used. In CHARMM-Drude simulations the charges from atomic cores, lone pairs, and Drude particles within a cutoff of 20 Å were all explicitly included. It is expected that the CHARMM-ECC scaled charges and CHARMM-Drude charges will significantly alter the local electrostatic environment of the SF carbonyls, compared to CHARMM simulations, and it is the main goal of this work to analyze the corresponding 2D IR spectra.

The nearest-neighbor frequency shifts (*ω_i,_*_nn_, NNFS) were determined using the approach of Jansen *et al.* [92, 93]. NNFS account for the frequency shifts due to covalent through-bond effects from the nearest-neighbor amide groups [92, 93]. These frequency shifts depend on Ramachandran angles as explained in Ref. 92.

The remaining contribution to the amide I frequency is the frequency shift due to ^13^C^18^O isotope labeling which was set to Δ*ω_i,_*_iso_ = *−*66 cm*^−^*^1^ based on previous experimental works [49, 50]. This shift is large enough to experimentally isolate peaks of the labeled carbonyls from the rest of the backbone carbonyls. In simulations, this energetic separation significantly reduces the computational cost because only isotope labeled amide I groups have to be included in simulations.

In general, amide I vibrations in proteins interact and the interactions modify their local frequencies and create couplings. Amide I modes in a protein can be coupled *via* through-bond and through-space [94]. In case of single-labeled residues in the SF which is what we model here, one residue in each of the four chains is labeled. In such case, because the isotope labeled amide I groups are on different KcsA chains, they interact through-space only. The coupling between them is described using the transition dipole coupling scheme [95]

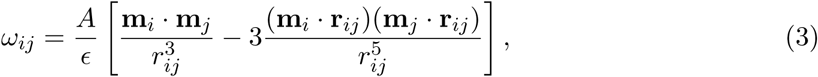

where **m***_i_* is the transition dipole of *i*th amide I mode (units D Å*^−^*^1^ *u^−^*^1^*^/^*^2^, *u* is the atomic mass unit), **r***_ij_* is the vector (in Å) connecting the transition dipoles of *i*th and *j*th amide I modes, and *r_ij_* is the corresponding distance. The dielectric constant is set to *ɛ* = 1 and the conversion factor *A* = 51.43 gives the coupling *ω_ij_* in the units of cm*^−^*^1^ [52]. The amide I transition dipole location, orientation, magnitude were derived from calculations by Torii and Tasumi [52, 95].

Based on the above description, the frequencies, couplings, and transition dipoles were calculated for every 20 fs snapshot along each 20 ps trajectory utilizing MDAnalysis library [96, 97]. Then the 2D IR spectra were calculated using the numerical integration of the Schrödinger equation (NISE) software [98–100] under parallel pump-probe polarization conditions and only *T*_2_ = 0 fs waiting time was considered. Previous 2D IR simulations and experiments on SF of KcsA showed that there are essentially experimentally discernible dynamics on picosecond timescales in the SF. Therefore no additional information can be obtained at longer waiting times. The vibrational anharmonicity was set to 14 cm*^−^*^1^ as determined experimentally in Ref. 101.

The simulation protocol described above was used previously in simulations of 2D IR spectra of ion channels using GROMOS96 [23, 24, 34, 53–55, 102] and amber99sb force fields [25, 56]. Cunha *et al.* [103] systematically compared linear and 2D IR amide I spectra simulated using various combinations of force fields, electrostatic frequency maps, and coupling models on a set of three small peptides. For each method they determined the frequency shifts that have to be applied to simulated spectra to maximize the overlap with experimental spectra. Note that Cunha *et al.* [103] used standard non-polarizable force fields CHARMM27, Amber99SB-ILDN, OPLS-AA, and Gromos-54a7, while this work focuses on the effects of including polarization on 2D IR spectra. The important for this work observation from Ref. [103] is that a good agreement between simulations and experiment can in some cases be achieved by simply shifting simulated spectra. However, as shown by Ryan *et al.* [24] 2D IR spectra of hardand soft-knock configurations differ not only by the frequency of the main peak but also by the number of peaks. Therefore, the simulated spectra presented below will be analyzed with a particular emphasis on the number and the origins of the peaks, since the peak frequency is already known to be sensitive to the force field.

## 3 Results and Discussion

Figure 2 shows 2D IR spectra of the KcsA channel simulated CHARMM, CHARMM-ECC, and CHARMM-Drude 2D IR spectra. Following experimental work [24, 35] the spectra are calculated with ^13^C^18^O isotope-labeled V76 and G77 residues. Experimental 2D IR spectra from Ref. [24] are shown as well. Note that both experiments and simulations show ^13^C^18^O isotope-labeled amide I peaks, but because the simulated peaks shift depending on many factors, as discussed above, the frequency region needed to show all the simulated peaks was extended compared to experimental spectra. The red dashed boxes in every simulated spectra show the corresponding experimental frequency ranges.

**Figure 2:**
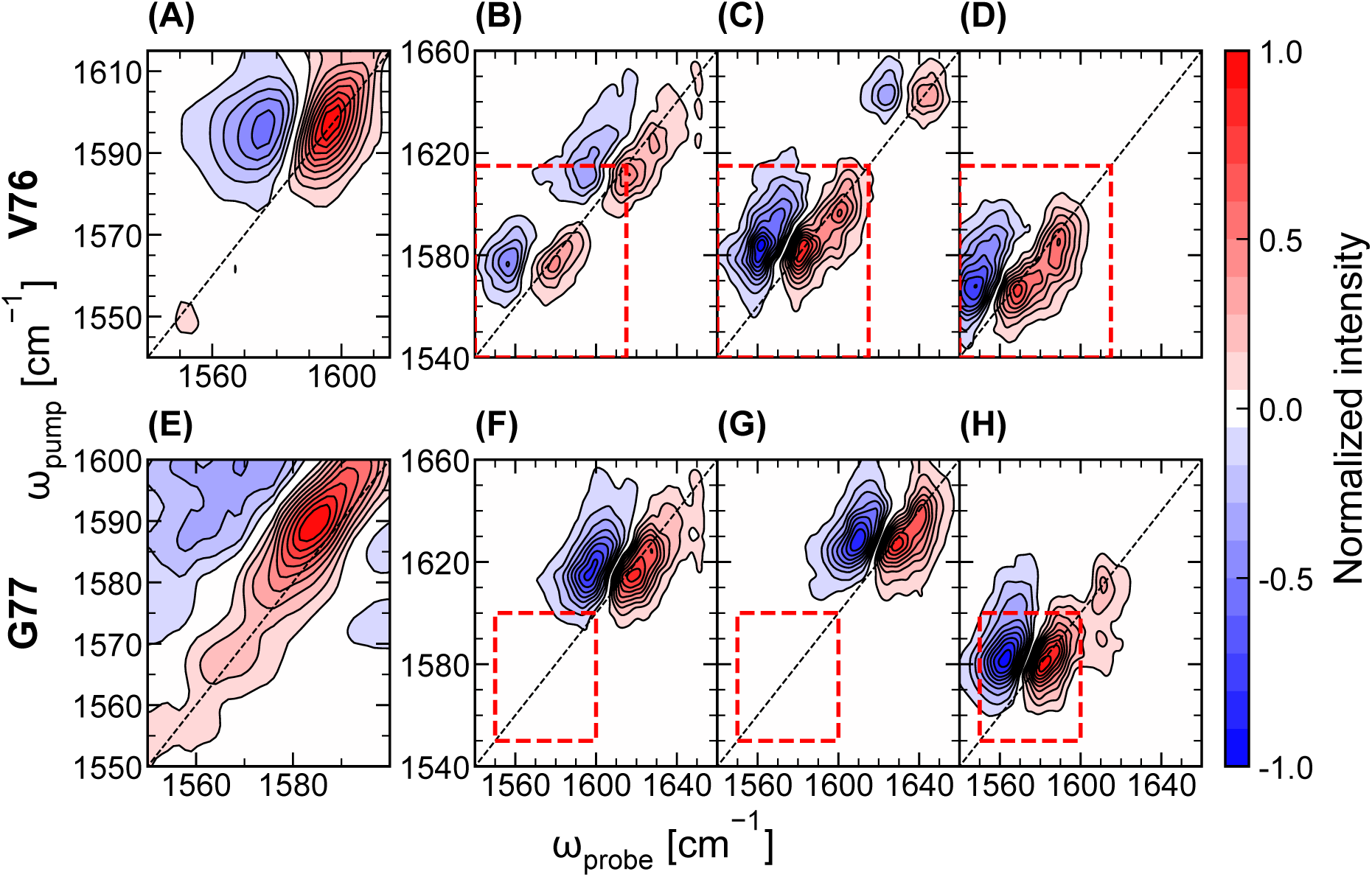
Top: Experimental (A) 2D IR spectrum of the ^13^C^18^O isotope labeled V76 residue and simulated 2D IR spectra using CHARMM36m (B), CHARMM-ECC (C), and CHARMM-Drude (D) force fields. Bottom: Experimental (E) 2D IR spectrum of the ^13^C^18^O isotope labeled G77 residue and simulated 2D IR spectra using CHARMM36m (F), CHARMM-ECC (G), and CHARMM-Drude (H) force fields. Experimental spectra are from Ref. [24]. The waiting time in all experimental 2D IR spectra is 500 fs. Simulated spectra correspond to 1:1 mixture of [KWKW] and [WKWK] configurations.

### 3.1 V76 spectra

As depicted in Figure 2A, the experimental 2D IR spectrum for isotope-labeled V76 carbonyl exhibits a pair of inhomogeneously broadened peaks centered at *ω*_pump_ = 1591 cm*^−^*^1^. The positive peak (red) arises from the 0*→*1 vibrational transition, while the negative peak corresponds to the 1*→*2 vibrational transition. The *ω*_probe_ frequency difference between the two peaks is the diagonal vibrational anharmonicity [39]. Experimentally no additional amide I peaks were found [24]. In contrast, the simulated 2D IR spectra contain a series of peaks for each force field studied. The V76 2D IR spectra from CHARMM force field, Figure 2B contains peaks at 1580 cm*^−^*^1^, 1616 cm*^−^*^1^, and less intense peaks at a higher frequency *ca.* 1629 cm*^−^*^1^. We focus on the two main features. To understand the origin of 1580 cm*^−^*^1^ and 1616 cm*^−^*^1^ peaks, in Figure 3A and B we plot the distribution of amide I frequencies calculated using Eq. 1 for each of the four protein sub-units and each configuration [KWKW] and [WKWK]. We observe that for the [KWKW] configuration all amide I frequencies are close and the peak is at 1575–1585 cm*^−^*^1^ which is directly attributed to the 1580 cm*^−^*^1^ peak in the total spectrum. It should be noted that because of the coupling between all amide I modes the resulting peaks shift from the uncoupled frequencies plotted in Figure 3, yet the latter are still useful in interpreting the spectra. The frequency distributions for the [WKWK] configuration is different from those of the [KWKW] configuration. Sub-units C and D produce nearly identical distributions both peaked at *ca.* 1600 cm*^−^*^1^. The frequency distribution from sub-unit B peaks at a lower frequency of 1575 cm*^−^*^1^. Interestingly, the sub-unit A produces a very broad distribution of frequencies with a peak at *ca.* 1600 cm*^−^*^1^ and a pronounced shoulder at larger frequencies. We attribute this shoulder to the configuration in which the V76 carbonyl flips away from the selectivity filter. Such a configuration was observed in previous MD simulations of KcsA and other K^+^ channels [23, 102, 104–112]. MD simulations suggest that the V76 flipped configuration has a pronounced impact on ion channel function including channel stability, ion selectivity, and C-type inactivation [105, 109, 113–116]. Experimental confirmation of the V76 flipped configuration is elusive [14, 114, 117]. When the V76 carbonyl flips away from the selectivity filter its electrostatic environment changes significantly resulting in increased vibrational frequency due to weaker electric fields behind the selectivity filter [23, 102]. The lower intensity of this peak is because V76 occurs in one out of four protein sub-units. Figure 4A shows the KcsA configurtion with one V76 carbonyl flipped.

**Figure 3:**
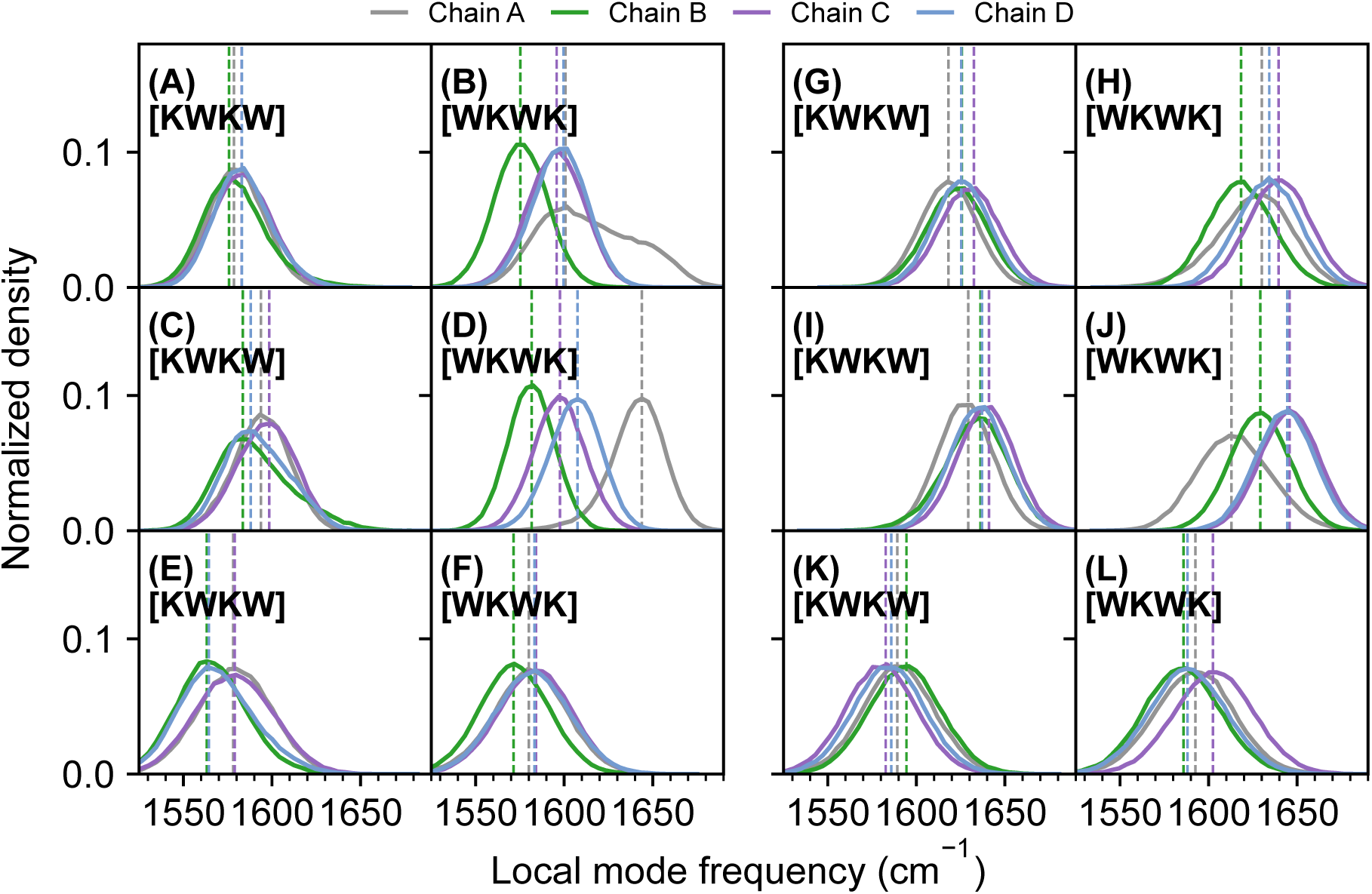
Distributions of the amide I local mode frequencies for V76 (A-F) and G77 (G-L) residues calculated separately for each of the four KcsA subunits (chains), two ion configurations, and all force fields studied here: CHARMM36m (A), (B), (G), and (H); CHARMM-ECC (C),(D), (I), (J); CHARMM-Drude (E), (F), (K), and (L).

**Figure 4:**
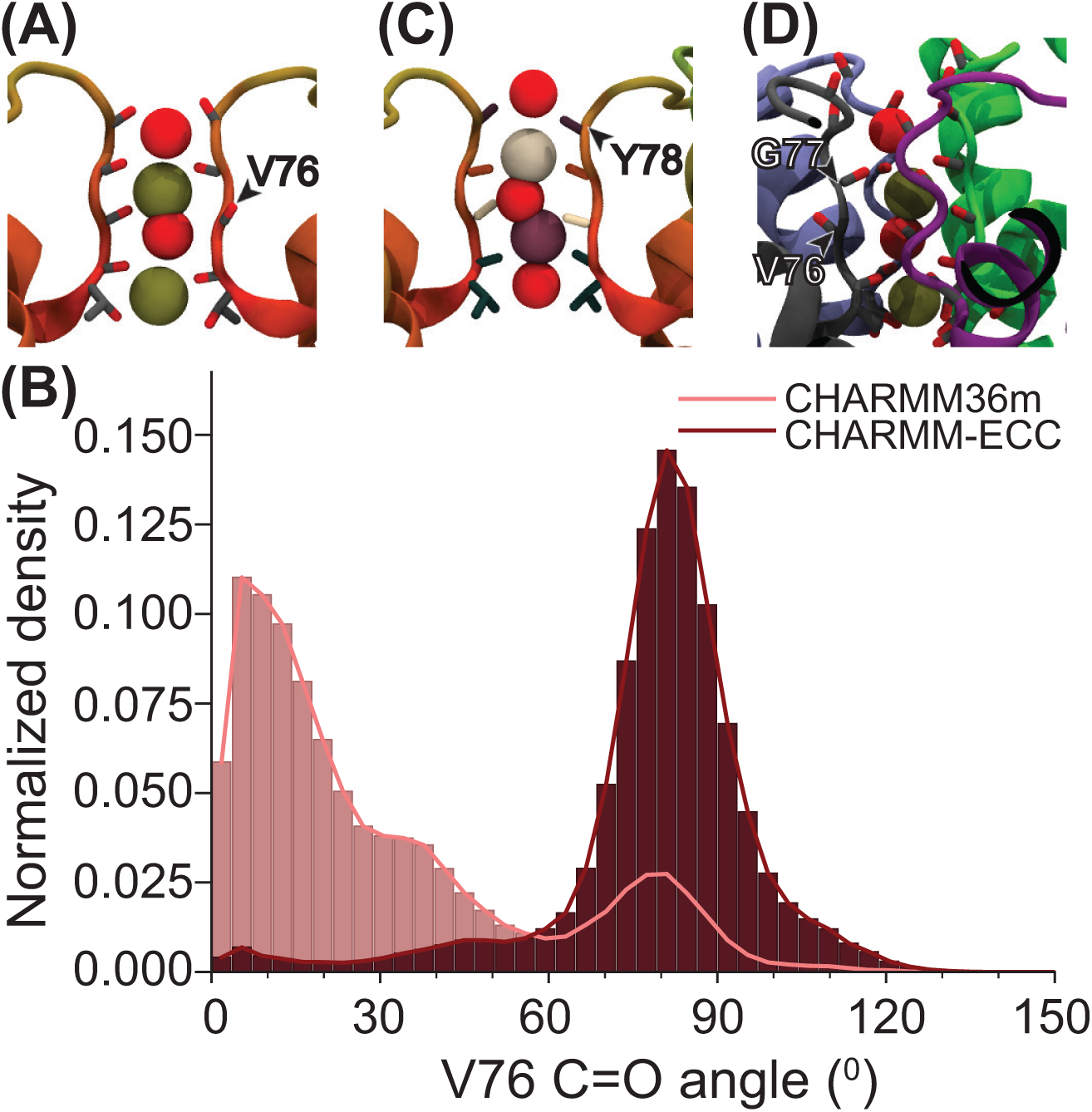
(A) A snapshot of the [WKWK] configuration with one of the V76 carbonyls flipped away from the pore. (B) Normalized distribution of the V76 C=O angle relative to the initial C=O angle calculated for the Chain A from CHARMM36m and CHARMM-ECC trajectories. Trajectories were pre-aligned using selectivity filter C*α* atoms (residues 75–79). (C) The snapshot of the [KWKW] configuration showing the ion in the S1 site (cream) slightly shifted towards the S2 site upon water molecule (red) entering the S0 site (from CHARMM36m simulations). (D) A snapshot of the [WKWK] configuration in CHARMM-ECC simulations, with the ion at the S2 site sticking closer to chains C and D due to V76 flipping disrupting ion coordination in chain A.

To summarize, previous simulations of 2D IR spectra of the isotope-labeled V76 carbonyl, based on GROMOS force field, showed that both [KWKW] and [WKWK] configurations yield peaks at *ca.* 1590 cm*^−^*^1^ which was integral in reproducing and interpreting the experimental spectra. Moreover, the V76 flipped configuration was found to be incompatible with 2D IR experiments [24, 34]. In contrast, CHARMM force field generates configurations that yield different frequencies leading to two distinct peaks and a third, lower intensity peak at 1629 cm*^−^*^1^ which results from the V76 flipped configuration. Approximately equal intensities of the two main peaks at 1580 cm*^−^*^1^ and 1616 cm*^−^*^1^ peaks are due to the equal ratio of [KWKW] and [WKWK] configurations used in simulations. Therefore, we conclude that non-polarizable CHARMM force field does not produce configurations of the SF that are compatible with 2D IR experiments and GROMOS is a better choice for a non-polarizable force field for this particular system.

Continuing the analysis of simulated V76 spectra we focus now on the CHARMM-ECC results. We note that, similar to CHARMM, CHARMM-ECC spectra also contain multiple peaks. The two main peaks appear at 1583 and 1600 cm*^−^*^1^, in a better agreement with experiment. We also observe minor peaks at 1643 cm*^−^*^1^, higher than the corresponding feature in the CHARMM spectra. Figures 3C and D show the distribution of amide I frequencies from CHARMM-ECC simulations. Multiple peaks, especially for the [WKWK] frequencies point to a broader variety of structural and electrostatic environments for amide I carbonyls in the CHARMM-ECC simulations compared to CHARMM (and CHARMM-Drude) MD simulations. The [KWKW] configuration contributes to the two major peaks in the total 2D IR spectrum, while the [WKWK] configuration, in addition, is responsible for a shoulder at *ca.* 1600 cm*^−^*^1^ and the 1643 cm*^−^*^1^ peak. The latter arises from V76 flipped configuration of the chain A which is the same as already identified in the CHARMM spectrum and discussed above. Figure 4B compares normalized distribution of the Chain A V76 C=O angle relative to the initial structure in CHARMM and CHARMM-ECC simulations. Trajectories were pre-aligned using selectivity filter C*α* atoms (residues 75–79). The three shoulders observed for CHARMM indicate various partial V76 carbonyl configurations which explain the broadening of the corresponding frequency distribution from Figure 3B. In contrast in the CHARMM-ECC simulations the V76 carbonyl only assumes a single 80*^◦^* partial flipped configuration. This explains a much narrower distribution of frequencies of the Chain A V76 carbonyl as shown in Figure 3D.

The high-frequency peak is completely absent in CHARMM-Drude 2D IR simulations of V76 2D IR spectrum shown in Figure 2D. The corresponding trajectory and frequency distribution (Figures 3E and F) show no V76 carbonyl flipping occurring. There are two main spectroscopic features, the main peaks 1569 cm*^−^*^1^ and a smaller intensity peaks at 1589 cm*^−^*^1^. To understand their origin we examine the corresponding frequency distributions shown in Figures 3E and F. We observe that both configurations [KWKW] and [WKWK] contribute to both spectral features.

To understand why the lower-frequency peak is more red-shifted in CHARMM-Drude spectrum (1569 cm*^−^*^1^) compared to CHARMM (1580 cm*^−^*^1^) and CHARMM-ECC (1583 cm*^−^*^1^), in Figure 5 we plot distributions of ion positions along the pore axis with the origin positioned at the geometric center of the selectivity filter C*α* atoms (residues 75–79) for [KWKW] and [WKWK] configurations. Focusing on the ion distributions near V76 (cream) we observe that for the [KWKW] configuration the K^+^ ions are positioned closer to the center of the S2 binding site, calculated here as a geometric average of the fluctuating positions of the V76 oxygen carbonyls. Closer ions create a stronger electric field causing larger frequency shifts. We conclude that the CHARMM-Drude simulations agree better with experiments due to the absence of the V76 flipped configurations.

**Figure 5:**
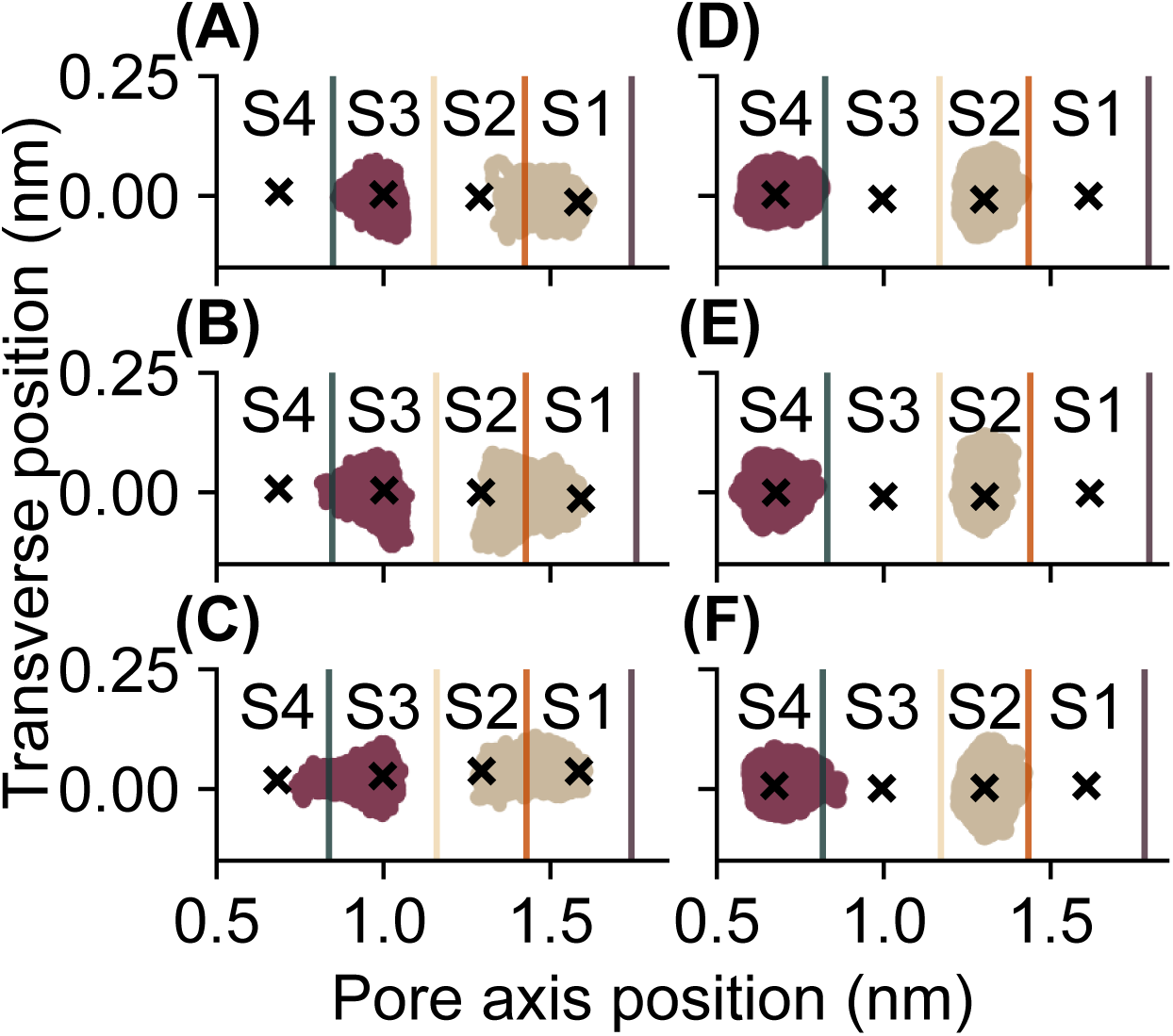
The distribution of ion positions along the pore axis with the origin positioned at the geometric center of the selectivity filter C*α* atoms (residues 75–79) calculated for [KWKW] (left) and [WKWK] (right) configurations from CHARMM36m (A and D), CHARMM-ECC (B and E), and CHARMM-Drude (C and F) trajectories. The “X” markers label to the ensembleaveraged geometric centers of each binding site. Vertical colored lines indicate the ensembleaveraged positions of the carbonyls: T75 (green), V76 (cream), G77 (orange), and Y78 (purple).

### 3.2 G77 spectra

The lower panel in Figure 2 compares experimental 2D IR spectra of ^13^C^18^O isotope-labeled G77 residues to the spectra simulated using CHARMM, CHARMM-ECC and CHARMM-Drude force fields. First, we note that the experimental spectrum contains one pair of peaks which is reproduced in all simulations, albeit the peak frequencies are different. The positive 0*→*1 (red) peak is more inhomogeneously broadened than the corresponding feature in the V76 spectrum. To quantify the extent of inhomogeneous broadening we use the ellipticity parameter (ELP) defined as [39]

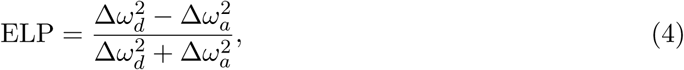

where Δ*ω_d_* and Δ*ω_a_* are the diagonal and antidiagonal linewidths, respectively. The linewidths were obtained by fitting Gaussian functions to the corresponding spectral profiles of positive peak. The ELP quantifies the relative elongation of a peak, with positive values indicating greater diagonal than antidiagonal elongation, zero indicating equal widths, and negative values indicating a greater antidiagonal width [39]. The calculated ELP values for the CHARMM, CHARMM-ECC, and CHARMM-Drude and experimental spectra are 0.55, 0.52, 0.45, and 0.50, respectively. All values of ELP values are within [0,1] interval, indicating noticeable diagonal elongation. In contrast, the ELP for the experimental V76 spectrum is 0.28.

The 2D IR spectrum from CHARMM simulations has a peak at 1618 cm*^−^*^1^ about 35 cm*^−^*^1^ blue-shifted with respect to the experimental spectrum. The distributions of the local amide I frequencies for [KWKW] and [WKWK] configurations shown in Figures 3G and H are very similar with the peaks in the 1620-1635 cm*^−^*^1^ frequency range which illustrates the inhomogeneous broadening of the spectral lineshapes.

CHARMM-ECC simulations produce an even more blue-shifted peak appearing at 1629 cm*^−^*^1^. In addition, there is a pronounced shoulder at the high-frequency side of the peak at 1641 cm*^−^*^1^. The ELP is 0.52 which is in a good agreement with the experimental value of 0.50. The amide I frequency distributions plotted in Figure 3I and J show very small differences for the [KWKW] configuration indicating that all G77 carbonyls in all protein subunits have very similar structural and electrostatic environments. In contrast, the G77 carbonyls of the KcsA sub-units of the [WKWK] configuration experience a broader variety of environments which is evidenced by the spread of the peaks of the frequency distributions from 1610 cm*^−^*^1^ (chain A) to 1645 cm*^−^*^1^ chains C and D. It is the amide I modes of these two chains that give rise to a shoulder in the simulated spectrum at 1641 cm*^−^*^1^. This change in local mode frequencies of G77 can be attributed to disrupted coordination at the S2 ion binding site as the flipped V76 carbonyl of chain A causes the ion to shift toward chains C and D.

The CHARMM-Drude G77 spectrum is in a very good agreement with experiment. The peak is located at 1582 cm*^−^*^1^ and the distributions of the local amide I mode frequencies point to similar environments for G77 carbonyls in all sub-units with an exception of the chain C in the [WKWK] configuration whose peak is at 1600 cm*^−^*^1^. The calculated ELP for the CHARMM-Drude spectrum 0.45 is also is a good agreement with the experimental value of 0.50.

The distribution of ion positions throughout the entire MD trajectory shown in Figure 5 explains the more significant inhomogeneous broadening observed in the G77 spectra compared to the V76 spectra. It is clear from Figure 5 that the ions sample a much broader range of distances to the carbonyl oxygen atoms compared to any other backbone carbonyl of the SF. Notably, ions initially placed at the center of S1 sites of the [KWKW] configuration were found to shift to the S2 site. The ion shifts from S1 to S2 binding sites are mediated by a water molecule entering the S0 site. The representative configuration from CHARMM simulation is shown in Figure 4B but we found the same water-mediated ion shifts to occur in simulations with all force fields studied here. Interestingly, we found that in the CHARMM-Drude simulation (Figure 5C) the shift of the S1 ion into the S2 site is accompanied by the shift of the from the S3 site to the S4 site which does not occur in CHARMM and CHARMM-ECC simulations.

## 4 Conclusion

Our simulations show that 2D IR spectra calculated based on MD trajectories are very sensitive to the treatment of the electronic polarizability. The best agreement with experimental spectra was achieved with Drude polarizability incorporated into CHARMM force field. The mean-field treatment of the polarizability within the ECC approach leads to minor improvements in the simulated spectra. Altogether, our simulations underscore the importance of explicit treatment of polarizability in MD simulations of ion channels. Our work is an example of an underexplored but promising approach to validating force fields based on 2D IR spectroscopy.

## Author Contributions

A.A.K. conceived the idea. A.A.K. and R.L.R. designed the research. R.L.R. carried out all simulations. A.A.K. and R.L.R. analyzed the data. A.A.K. and R.L.R. wrote the manuscript.

## Acknowledgments

Research reported in this publication was in part supported by the National Institute of General Medical Sciences of the National Institutes of Health under Award R35GM150963. The content is solely the responsibility of the authors and does not necessarily represent the official views of the National Institutes of Health. A.A.K. acknowledges the support from the start-up funds provided by the College of Arts and Sciences and the Department of Physics and Astronomy of the University of Delaware. Calculations were performed with high-performance computing resources provided by the University of Delaware.

## Notes

### Competing Interest Statement

The authors have declared no competing interest.

